# Mapping vegetation types in Antarctic Peninsula and South Shetlands islands using Sentinel-2 images and Google Earth Engine cloud computing

**DOI:** 10.1101/2021.09.14.460232

**Authors:** Eliana Lima da Fonseca, Edvan Casagrande dos Santos, Anderson Ribeiro de Figueiredo, Jefferson Cardia Simões

## Abstract

The Antarctic vegetation maps are usually made using very high-resolution images collected by orbital sensors or unmanned aerial vehicles, generating isolated maps with information valid only for the time of image acquisition. In the context of global environmental change, mapping the current Antarctic vegetation distribution on a regular basis is necessary for a better understanding of the changes in this fragile environment. This work aimed to generate validated vegetation maps for the North Antarctic Peninsula and South Shetlands Islands based on Sentinel-2 images using cloud processing. Sentinel-2 imagery level 1C, acquired between 2016 and 2021 (January-April), were used. Land pixels were masked with the minimum value composite image for the “water vapor” band. The NDVI maximum value composite image was sliced, and its classes were associated with the occurrence of algae (0.15 – 0.20), lichens (0.20 – 0.50), and mosses (0.50 – 0.80). The vegetation map was validated by comparing it with those from the literature. The present study showed that Sentinel-2 images allow building a vegetation type distribution map for Antarctica Peninsula and South Shetlands Islands.

## 1. Introduction

The vegetation in the Antarctic environment is restricted to ice-free areas, mainly in the Antarctic islands and in the coastal areas of the continent regions (Alberdi et al., 2002; Convey, 2006; Fretwell et al., 2011). These plant communities are predominantly cryptogamic, also known as lower plants or biological soil crusts (BSC) (Convey, 2010), and their growth season length depends on the climatic conditions, latitude, and relief (Selkirk & Skotnicki, 2007). The availability of liquid water is the most critical factor for the development of vegetation communities in Antarctica, which is available during few months when snow melts and summers rain occurs, or when the humidity can be absorbed directly from the air (Elster, 2002; Bölter et al., 2002; Choi et al., 2015). As a result, the expansion of vegetated areas occurs at a very slow rate in the Antarctic Maritime region (Fritsen & Priscu, 1998; Convey, 2006).

The use of remote sensing data to map Antarctic vegetation is concentrated in areas frequently visited by researchers (Calviño-Cancela & Martin-Herrero, 2016), usually made at local scales, with very-high-resolution images collected by orbital sensors, such as KOMPSAT-2 and QuickBird (Shin et al., 2014) and WorldView-2 (Jawak et al., 2019) and with unmanned aerial vehicles (UAV) (Miranda et al., 2020; Sotille et al.,2020). These studies primarily focused on detecting the presence/absence of vegetation, generating maps with information valid only for the time of image acquisition. At regional scales, based on Landsat Image Mosaic of Antarctica (LIMA mosaic) generated using images acquired between 2007/2008, Fretwell et al. (2011) mapped the probability of vegetation occurrence for the Antarctic Peninsula.

In the context of global environmental change, mapping the current Antarctic vegetation distribution is required to understand the impact of these changes on vegetation biomass accumulation and the greenhouse gas cycle in the near future. Even in recent literature, the Antarctic vegetation is not considered in global forecasts (e.g., Jung et al., 2021) due to the lack of information over this region. A valid vegetation map can be used as an input layer in simulations processes about environmental changes, besides, it is analyzed by itself content. This work aimed to generate a vegetation map for the northern Antarctic Peninsula and South Shetlands based on Sentinel-2 images using Google Earth Engine (GEE) data catalog and cloud processing (Gorelick et al., 2017).

## 2. Methods

All the processes were made in GEE using its native JavaScript interface, accessed by Google Chrome. The cartography was made using QGIS and ESRI satellite basemap, available at Quickmapservices plugin. To generate the vegetation maps eleven subsets were created, being four for the northern Antarctic Peninsula (south peninsula, north peninsula, north islands, west islands) and seven for the South Shetlands Islands (Elephant and Clarence, King George, Nelson, Robert, Livingston, Deception, Smith, and Low), in order to not exceed the processing capacity quota to calculate the total mapped area. Monthly meteorological data, from ECMWF/ERA5 climate reanalysis, such as total precipitation, mean air temperature, and total net shortwave radiation (presented as supplementary material), were used to analyze the differences between mapped areas over different subsets.

### 2.1 Ocean/land mask generation

For automatic vegetation mapping, mask the ocean areas is required once the phytoplankton at the ocean also made photosynthesis, and can be mapped as vegetation over the ocean areas, as observed by Fretwell et al. (2011). The vector edges from the Antarctic Digital Database, available at Quantarctica packaged (Matsuoka et al., 2021), were generated using a small-scale, so the edges have low detail levels (as can be observed at supplementary material) and can not be used to mask land areas.

An ocean/land mask was built over the area using the “water vapor band” (B9), centered at 945nm and 943.2nm for Sentinel 2A and 2B, with 60 meters spatial resolution. At these wavelengths, the water reflectance is null, allowing the separation of water and non-water pixels while, for the same wavelengths, the cloud and snow reflectance is around 0.7 (Jansen, 2000). Using all the available images since January 1, 2016, a minimum value composite image for B9 was computed for the entire area by selecting its minimum value registered for each pixel for the whole time series. With this approach, every water pixel acquired without ice cover, cloud cover, or phytoplankton, at least once during the time series, will be filled with TOA reflectance value near zero, due to its spectral reflectance pattern. The B9 minimum value composite image had its values remapped, building a binary image used as ocean/land mask, labeling those pixels with values lower than 120 as water and all others pixels as non-water.

### 2.2 Vegetation type identification

The vegetation type identification was made using a Sentinel-2 image, acquired on February 23, 2019, from Surface Reflectance Sentinel-2 collection (COPERNICUS/S2_SR), available at the GEE data catalog. The option of using surface reflectance data was due to the availability of information from the literature that considers surface reflectance data to BSC target identification. Also, the availability of a Sentinel-2 cloud-free image acquired over Harmony Point (Figure 2a) for the same month when the fieldwork was made was considered. The fieldwork was carried out on February 13 - 20, 2015, when information about vegetation cover type, i.e., algae, lichens, or mosses (Figure 3) was collected over 17 samples points in the Harmony Point. For labeled the NDVI classes, the surface reflectance profiles from Sentinel-2 bands located at blue, green, red, red-edge, NIR, and SWIR wavelengths were generated for each NDVI interval (for every 0.05) and compared with literature (Lovelock & Robinson, 2002; Zhang et al., 2007) and with fieldwork information.

**Figure 1.**
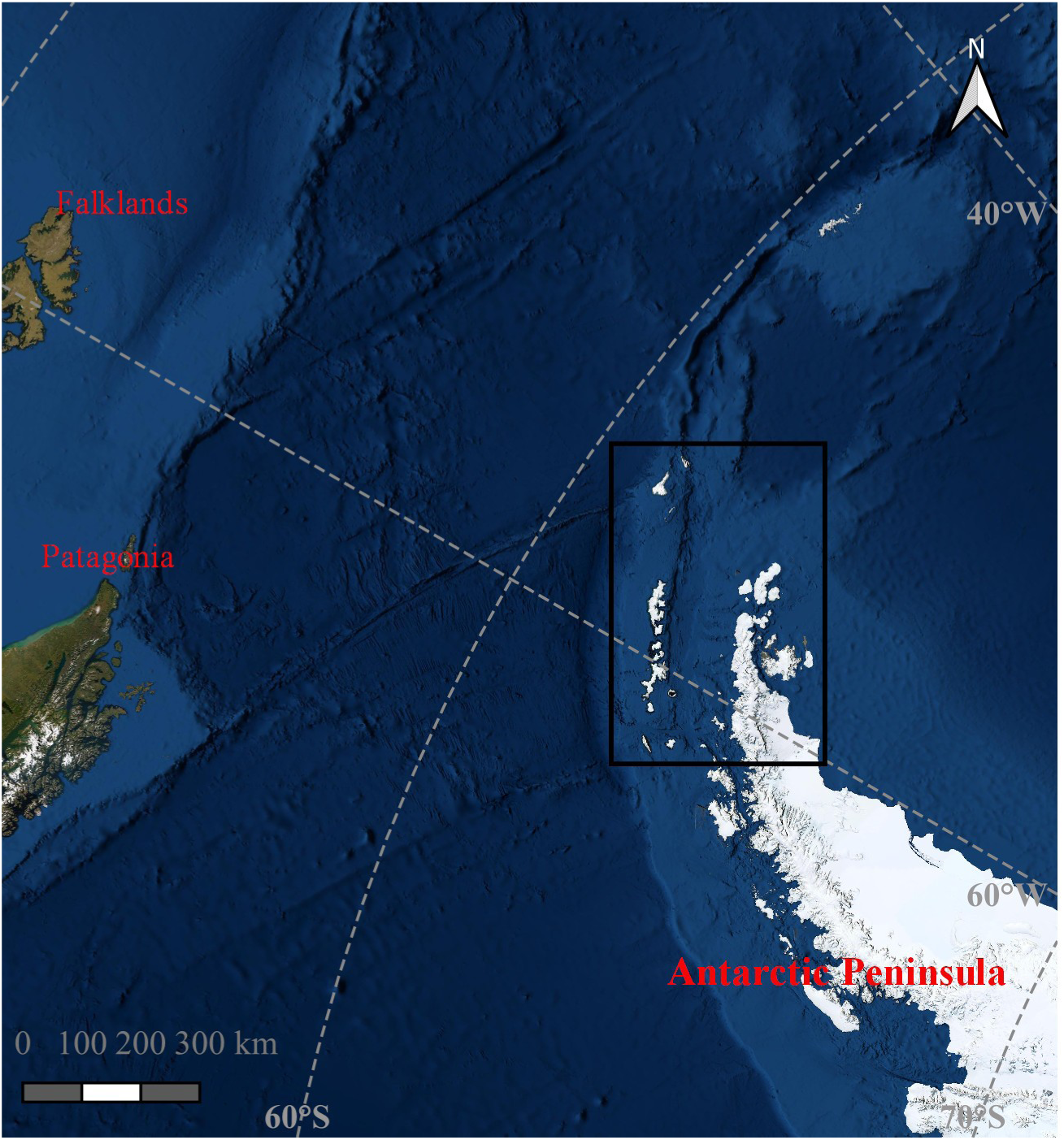
Northern Antarctic Peninsula and South Shetlands (black rectangle) over ESRI satellite basemap.

**Figure 2.**
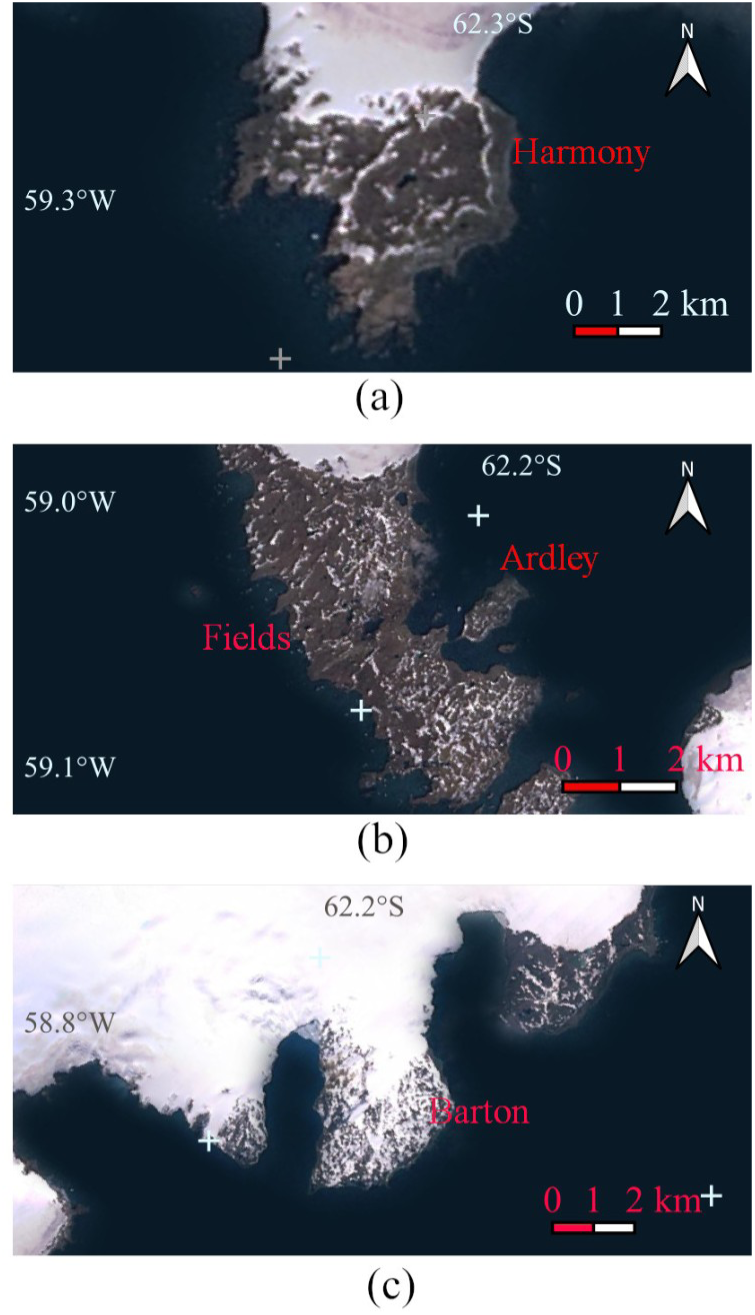
Harmony Point (a), Fildes Peninsula and Ardley Island (b), and Barton Peninsula (c) over ESRI satellite basemap.

**Figure 3.**
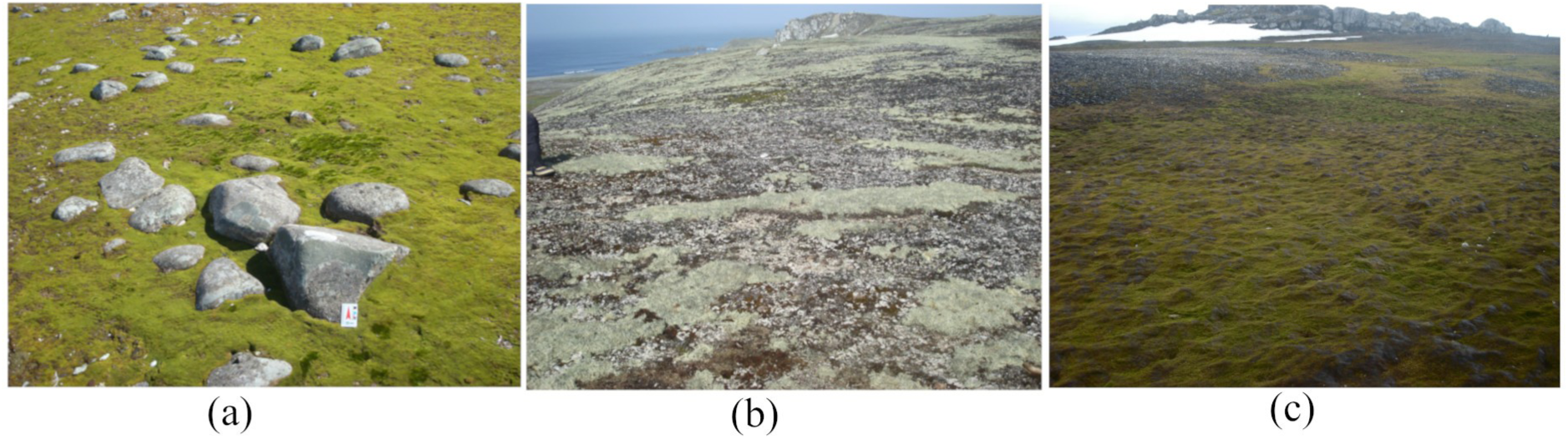
Biological soil crusts at Harmony point: algae (a), lichens (b), mosses (c)

### 2.3 Vegetation maps from NDVI images

The dataset was composed of all Sentinel-2 images at level 1C (orthorectified and radiometrically corrected at the top-of-atmosphere (TOA) reflectance), available at GEE, (“COPERNICUS/S2” collection), acquired by Sentinel 2A and 2B over the northern portion of the Antarctic Peninsula and its surrounding islands and South Shetlands islands from 2016 until 2021. These images contain the TOA reflectance values calculated for each Sentinel-2 spectral band and quality assessment (QA) bands. The TOA reflectance product was used instead of the surface reflectance product due to the high spectral absorption and scattering of ocean optical constituents (IOCCG, 2020), that making with the atmospheric correction models in coastal areas did not work properly (Warren et al., 2019).

The NDVI images were calculated using Sentinel-2 bands located at red and near infra-red wavelengths (B4 and B8), with 10 meters spatial resolution. The Sentinel-2 cloud-mask and ice-snow mask filters cannot be appropriately applied over this region, whether built using QA60 band or from QA10 and QA20 bands. When these masks were applied over a Sentinel-2 image acquired over the Antarctic Peninsula and South Shetlands, it masked all image pixels, returning only empty pixels instead of the TOA reflectance values.

To avoid the influence of cloud and snow pixels on NDVI values, a unique maximum value composite image was computed using all the available images collected over the study area from January 1 to April 30, which are the summer/autumn months in the Southern hemisphere, and are related to the vegetation growth season in the Antarctic environment (Alberdi et al., 2002; Elster, 2002; Lewis-Smith, 2007; Selkirk & Skotnicki, 2007). This approach is based on NDVI proprieties, and its values range, between -1 and 0, for water, ice, clouds, and clouds shadows and between 0 and 1 for bare areas and vegetation. Also, a NDVI maximum value composite image was computed for each analyzed year to generate vegetation maps on an annual basis. Each NDVI maximum value composite image had its values sliced into 21 classes, the first class for the negative values and the positive values were sliced for each 0.05 until 1. The NDVI classes were labeled as mosses, lichens or algae, for generating the vegetation map. The vegetation maps were validated by comparing with results from Andrade et al. (2018) and Sun et al. (2021) over Fildes Peninsula and Ardley Island (Figure 2b) and Shin et al. (2014) over Barton Peninsula (Figure 2c).

## 3. Results

### 3.1 Vegetation class labels

Over the Harmony point, the NDVI values associated with vegetated areas ranged from 0.15 to 0.8, being similar to the observed by Fretwell et al. (2011), who associated NDVI values higher than 0.20 with vegetation pixels in Antarctic Peninsula using Landsat images. The surface reflectance values for the optical bands at blue, green, red, red edge, NIR, and SWIR wavelengths collected over each NDVI class are given in Table 1. The vegetation pixels were labeled as algae when NDVI values were between 0.15 – 0.20, lichens for 0.20 – 0.50, and mosses for 0.50 – 0.80, defined based on samples location and on surface reflectance patterns for each NDVI class. Algae presented lower NDVI values (0.15 – 0.20) than other BSC, being these values similar as observed by Yun et al. (2017). The increase in reflectance values at the red edge and NIR wavelengths (Table 1) did the criteria for separating mosses and lichens, based on results from Lovelock & Robinson (2002) and Zhang et al. (2007). Five sample points were collected in areas with BSC mosaic cover, and, in fact, were located in a midlle of two NDVI classes border and didńt used to label NDVI classes.

**Table 1.** Mean surface reflectance values for each spectral band collected over a Sentinel-2 image acquired on February 23, 2019, for each NDVI class at Harmony Point.

| NDVI class<br>interval | 0.426-0.558 /<br>B2 /<br>Blue | 0.523-0.595 /<br>B3 /<br>Green | 0.634-0.696 /<br>B4 /<br>Red | 0.688-0.720 /<br>B5 /<br>Red<br>Edge | 0.724-0.754 /<br>B6 /<br>Red<br>Edge | 0.760-0.800 /<br>B7 /<br>Red<br>Edge | 0.727-0.939 /<br>B8 /<br>NIR | 0.842-0.886 /<br>B8A /<br>NIR | 1.516-1.705 /<br>B11 /<br>SWIR | 2.001-2.371 /<br>B12 /<br>SWIR |
| --- | --- | --- | --- | --- | --- | --- | --- | --- | --- | --- |
| 0.15 - 0.20 | 0.12 | 0.14 | 0.14 | 0.16 | 0.18 | 0.18 | 0.20 | 0.19 | 0.21 | 0.16 |
| 0.20 - 0.25 | 0.08 | 0.10 | 0.11 | 0.13 | 0.15 | 0.16 | 0.18 | 0.18 | 0.25 | 0.17 |
| 0.25 - 0.30 | 0.07 | 0.09 | 0.10 | 0.13 | 0.15 | 0.17 | 0.19 | 0.19 | 0.27 | 0.18 |
| 0.30 - 0.35 | 0.07 | 0.09 | 0.10 | 0.13 | 0.16 | 0.18 | 0.20 | 0.20 | 0.27 | 0.18 |
| 0.35 - 0.40 | 0.07 | 0.09 | 0.10 | 0.13 | 0.17 | 0.19 | 0.21 | 0.21 | 0.29 | 0.19 |
| 0.40 - 0.45 | 0.06 | 0.09 | 0.10 | 0.13 | 0.17 | 0.20 | 0.22 | 0.22 | 0.29 | 0.19 |
| 0.45 - 0.50 | 0.06 | 0.08 | 0.09 | 0.13 | 0.18 | 0.21 | 0.24 | 0.24 | 0.31 | 0.20 |
| 0.50 - 0.55 | 0.05 | 0.08 | 0.09 | 0.13 | 0.19 | 0.22 | 0.25 | 0.25 | 0.31 | 0.20 |
| 0.55 - 0.60 | 0.05 | 0.08 | 0.08 | 0.13 | 0.20 | 0.24 | 0.28 | 0.28 | 0.32 | 0.20 |
| 0.60 - 0.65 | 0.04 | 0.07 | 0.07 | 0.13 | 0.22 | 0.25 | 0.30 | 0.30 | 0.33 | 0.20 |
| 0.65 - 0.70 | 0.04 | 0.07 | 0.07 | 0.14 | 0.25 | 0.30 | 0.36 | 0.36 | 0.33 | 0.20 |
| 0.70 - 0.75 | 0.03 | 0.07 | 0.06 | 0.14 | 0.27 | 0.32 | 0.38 | 0.38 | 0.32 | 0.19 |
| 0.75 - 0.80 | 0.03 | 0.06 | 0.06 | 0.13 | 0.26 | 0.32 | 0.37 | 0.38 | 0.29 | 0.17 |

### 3.2 Mapped vegetation areas

The final vegetation map at its full spatial resolution can be accessed by running the script inside the GEE platform, which the link is available as supplementary material. The mapped areas for the final vegetation map and for all analyzed years are presented in Tables 2 and 3. Over Antarctic Peninsula (Table 2) 155.7 km^2^of vegetation areas were mapped, and the algae are the most abundant vegetation type for all subsets. Over South Shetlands (Table 3), 60.4 km^2^ of vegetation areas were mapped, and the lichens were the most abundant vegetation type among all subsets.

**Table 2.** Mapped vegetation areas (km^2^) for algae (AL), lichens (LI), and mosses (MO) over the Antarctic Peninsula subsets for the study period (2016-2021) and for the final vegetation map.

| Subset ID* | NP_1 |  |  | NP_2 |  |  | NP_3 |  |  | NP_4 |  |  |
| --- | --- | --- | --- | --- | --- | --- | --- | --- | --- | --- | --- | --- |
| Vegetation type | AL | LI | MO | AL | LI | MO | AL | LI | MO | AL | LI | MO |
| 2016 | 2.80 | 2.00 | 0.02 | 1.32 | 1.23 | 0.07 | 4.81 | 6.58 | 0.18 | 11.60 | 7.39 | 0.74 |
| 2017 | 0.57 | 0.24 | 0.00 | 0.85 | 0.44 | 0.00 | 3.11 | 4.29 | 0.02 | 3.06 | 0.97 | 0.00 |
| 2018 | 2.51 | 1.29 | 0.00 | 1.53 | 0.57 | 0.00 | 3.10 | 4.01 | 0.03 | 8.81 | 3.22 | 0.00 |
| 2019 | 1.48 | 2.17 | 0.01 | 1.24 | 0.86 | 0.00 | 4.47 | 4.85 | 0.16 | 8.23 | 4.82 | 0.07 |
| 2020 | 0.45 | 0.30 | 0.00 | 1.55 | 0.70 | 0.00 | 5.69 | 4.40 | 0.06 | 18.29 | 8.08 | 0.02 |
| 2021 | 2.96 | 1.72 | 0.01 | 3.81 | 2.14 | 0.01 | 9.44 | 5.80 | 0.01 | 30.99 | 18.52 | 0.08 |
| Final map | 9.26 | 6.21 | 0.04 | 8.37 | 4.51 | 0.08 | 16.15 | 12.54 | 0.40 | 64.81 | 32.43 | 0.88 |

\* NP\_1 south peninsula, NP\_2 north peninsula, NP\_3 north islands, NP\_4 west islands

**Table 3.**
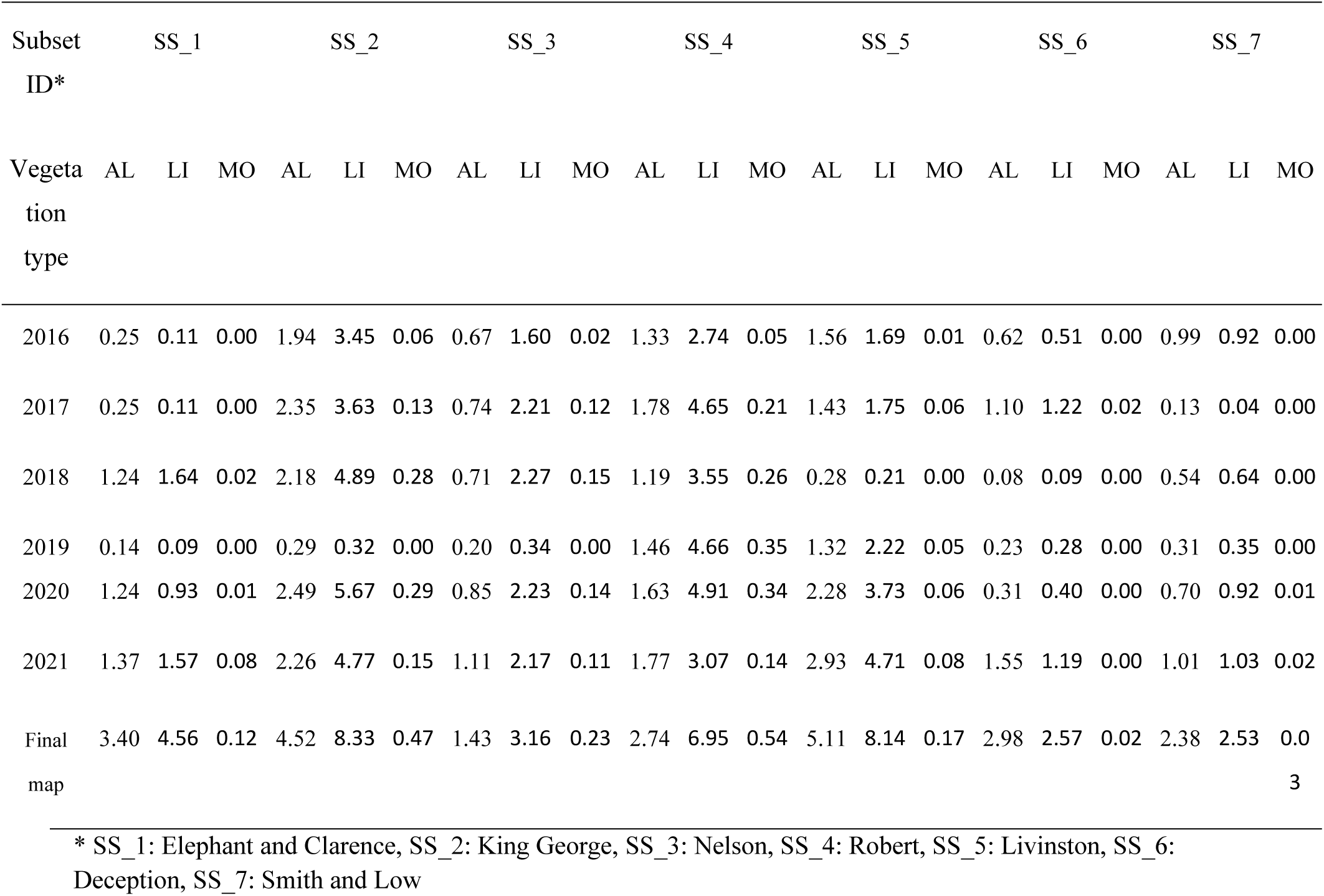
Mapped vegetation areas (km^2^) for algae (AL), lichens (LI), and mosses (MO) over South Shetlands islands subsets for the study period (2016-2021) and for the final vegetation map.

## 4. Discussion

Despite the availability of a long-term Landsat image series collected over the Antarctic at official datasets repositories, those images also do not have a correct georeferencing over the Antarctic, being the reason for the nonexistence of vegetation map over this region on a regular basis. The Sentinel-2 imagery is georeferenced using the Copernicus Precise Orbit Determination (POD), allowing georeferencing images based on satellite position at the acquired time. Thus, it solves the first fundamental problem of composing a satellite image time-series that is the correct georeferencing, which is essential for automatic data processing. The NDVI maximum value composite approach, in association with the ocean/land mask, was able to eliminate the cloud and cloud-shadows pixels and excluding those pixels from phytoplankton over the ocean, allowing the continuity of automatic processes.

The subsets have different area dimensions and, hence, a direct quantitative comparison of the maps is not possible. However, a high vegetation area mapped at subset NP_4 locate over the west islands can be noticed (Table 2), similar to results from Fretwell et al. (2011) over the same region. Furthermore, it can be observed that a great total net shortwave radiation amount at subset NP_4 in relation with other subsets (see at supplementary material), evidencing a microenvironment over this region that provided vegetation growth conditions. Zhou et al. (2021) also detected these microenvironment effects using remote sensing techniques to monitor the environmental changes, such as dry-snow line variations. Also, they were pointed out as one of the driving forces to other environmental changes (e.g., changes in phytoplankton communities) (Ferreira et al., 2020). Over the South Shetlands Islands, no relationships among vegetation distribution and the weather variables or the geographic location were found.

The annual variations in mapped vegetation areas observed among analyzed years (Tables 2 and 3) cannot be considered as a land cover change in this region. As pointed out by Shin et al. (2014), some variations in vegetation abundance in relation with the month of data acquisition and interannual meteorological conditions are expected, but not for the vegetation distribution area, since the expansion of vegetated areas occurs at a very slow rate in the Antarctic (Fritsen & Priscu, 1998; Convey, 2006). The differences in mapped vegetation areas among years can be observed in Figure 4, where the annual maps over Fildes Peninsula and Ardley Island are presented. Since no weather variations among the years were found over this area, these variations observed are, in fact, due to the cloud cover, which justifies the use of images acquired in more than one year to build a valid vegetation map from Sentinel-2 images.

**Figure 4.**
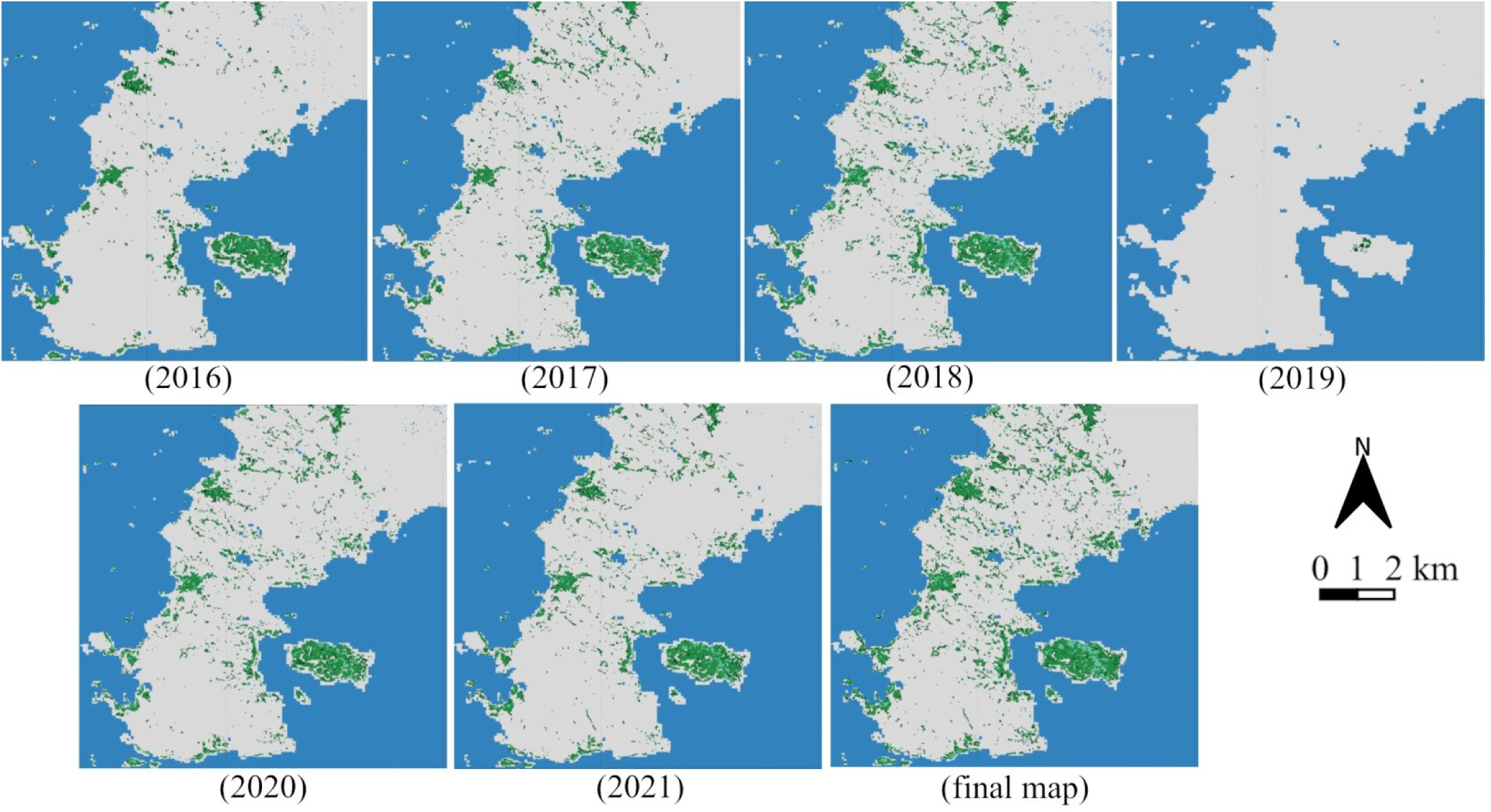
The annual vegetation distribution maps over Fildes Peninsula and Ardley Island for the study period (2016-2021) and for the final vegetation map.

### 4.1. Vegetation map validation

Over the Fildes Peninsula and Ardley Island, the final vegetation map (Figure 5a) were compared with results from Sun et al. (2021), who estimated the areas dominated by mosses and lichens using spectral mixture analysis with a WorldView-2 image and field measurements and with results from Andrade et al. (2018) who mapped the vegetation using a QuickBird image. Over Ardley Island, an area of 0.106 km^2^was estimated as occupied by algae, 0.659 km^2^ by lichens, and 0.212 km^2^by mosses. Sun et al. (2021) mapped lichens and mosses and estimated 0.3259 km^2^covered by mosses. Andrade et al. (2018) mapped two different classes, one for mosses (0.3165 km^2^) and the other for lichens and mosses association (0.3804 km^2^). Some differences in mapped areas are expected due to the spatial resolution of the images used. Note that, in many cases, the mosses distribution estimated with WorldView-2 (Sun et al., 2021) and QuickBird (Andrade et al., 2018) were mapped as algae using Sentinel-2 images. Both mosses and algae occur at the same moist microenvironment (Becker, 1982; Broday, 1996; Lovelock & Robinson, 2002), which can explain the greater difference observed in mapped area with mosses between this work and the areas mapped by Andrade et al. (2018) and Sun et al. (2021).

**Figure 5.**
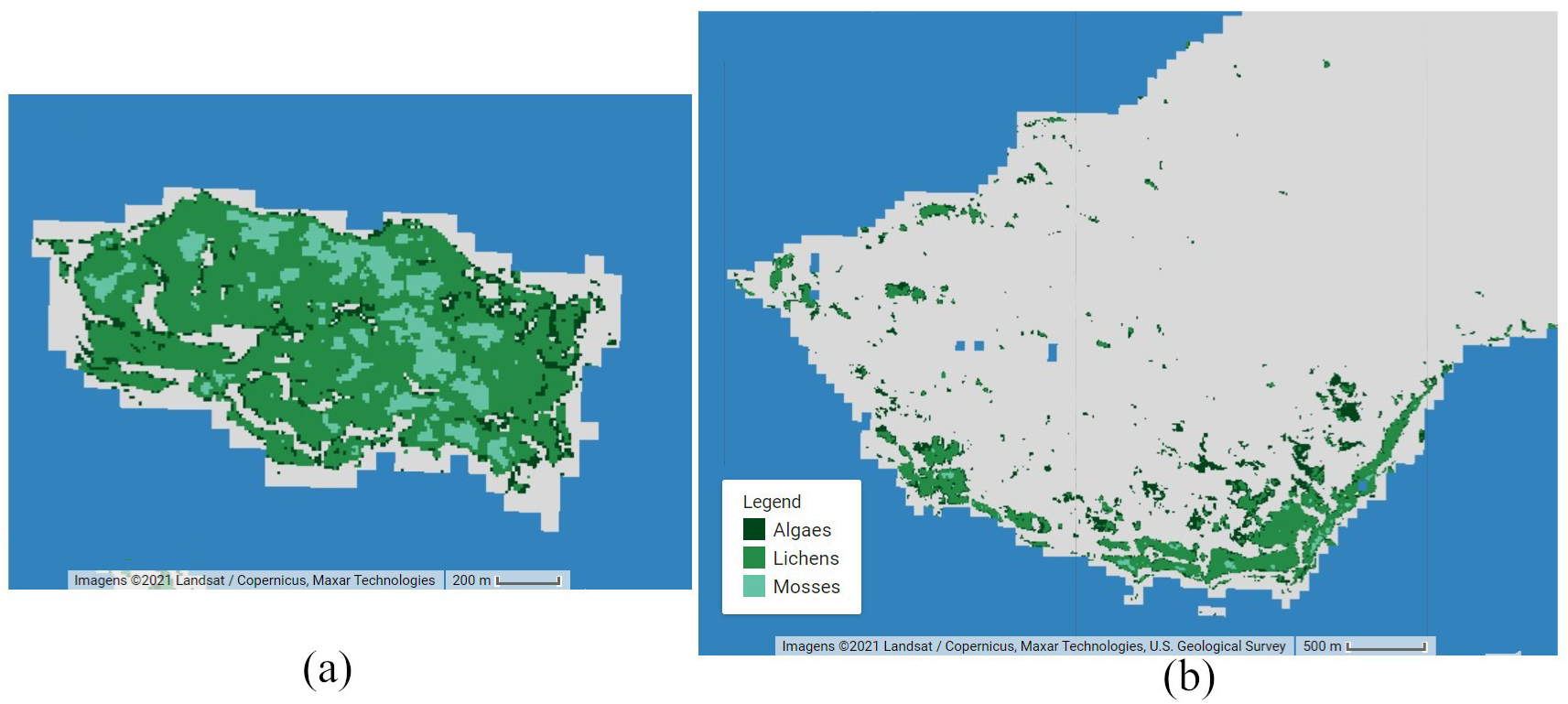
Vegetation map for Ardley Island (a) and for Barton Peninsula (b)

The most relevant comparison that can be made is about vegetation spatial distribution found over Fildes Peninsula and Ardley Island that was the same in all three works, indicates that our approach to map vegetation using Sentinel-2 images with 10 meters spatial resolution (B4 and B8) are valid with good results, similar to that obtained with very-high spatial resolution images. Over the Barton Peninsula, Shin et al. (2014) mapped vegetation abundance using KOMPSAT-2 and QuickBird very-high resolution images, by spectral mixture analysis and the spatial distribution of vegetation found over this area by these authors was similar to the map generated in the present study (Figure 5b), which indicates, again, that the approach using Sentinel-2 images was able to map the vegetation over the Antarctic environment. The absence of information with correct geo-location over the Antarctic region made validating the results a complex task. Since the mapped vegetation distribution found in this work was similar to that of Shin et al. (2014), Andrade et al. (2018), and Sun et al. (2021), the NDVI range from 0.15 – 0.80 can be considered valid to map the vegetation cover over the Antarctic Peninsula and South Shetlands islands using Sentinel-2 images.

## Author contributions

ELF performed the fieldwork, analyzed the data, wrote, reviewed and edited the manuscript; ECS performed the literature review, analyzed the data and wrote the manuscript; ARF performed the fieldwork; JCS reviewed the manuscript and acquired the financial resources of this research. All authors discussed the results and approved the final version of the manuscript.

